# Perineuronal Nets in the Insula Regulate Aversion-Resistant Alcohol Drinking

**DOI:** 10.1101/504571

**Authors:** H. Chen, A. W. Lasek

## Abstract

One of the most pernicious characteristics of alcohol use disorder is the compulsion to drink despite negative consequences. The insular cortex (insula) controls decision-making under conditions of risk or conflict and regulates maladaptive behaviors in the context of addiction. Cortical activity is tightly controlled by fast-spiking inhibitory interneurons that are often enclosed by specialized extracellular matrix structures known as perineuronal nets, which regulate neuronal excitability and plasticity. Using a mouse model of compulsive drinking in which alcohol was adulterated with the bitter tastant quinine, we demonstrate that disrupting perineuronal nets in the insula rendered mice more sensitive to quinine-adulterated alcohol. Activation of the insula, as measured by c-fos expression, occurred during aversion-resistant drinking and was further enhanced by elimination of perineuronal nets. These results provide fundamental insight into neuroanatomical and cellular mechanisms that control compulsive drinking.

## Introduction

Alcohol use disorder (AUD) is a common psychiatric disorder with an estimated lifetime prevalence of 29% (*1*). AUD is a chronic relapsing condition characterized by loss of control in limiting alcohol intake, a negative emotional state when alcohol use is discontinued, and compulsive drinking. Compulsive drinking, defined as the pathological motivation to consume alcohol, manifests as drinking despite negative consequences, such as job loss, legal problems, and damage to interpersonal relationships. This type of risky drinking is not addressed by currently approved AUD pharmacotherapies (*2*). The identification of the underlying brain mechanisms that promote compulsive drinking is critical for effective development of new interventions to treat AUD.

One brain area that regulates compulsive drinking is the anterior insula (*3, 4*), a cortical region located deep within the lateral sulcus (*5*). The importance of the anterior insula in addiction emerged just over a decade ago, when Naqvi *et al* demonstrated that smokers who had stroke damage in the insula were more likely to quit smoking and reported less craving for cigarettes (*6*). More recent studies have demonstrated that alcohol-dependent subjects have decreased anterior insula volume and cortical thinning compared with control subjects and these structural deficits are correlated with higher compulsive drinking measures (*7*). The anterior insula is also activated during risky-decision making in individuals with AUD, with decreased activation associated with increased risk-taking (*8*).

Compulsive drinking is modeled in animals by pairing alcohol seeking or intake with an aversive consequence such as an electric foot shock or addition of bitter-tasting quinine to the alcohol solution (*9*). Compulsive animals are aversion-resistant, meaning that they willingly seek and consume alcohol despite the threat of shock or aversive taste. Optogenetic inhibition of excitatory inputs from the insula to the nucleus accumbens reduced aversion-resistant alcohol intake in rats (*4*), demonstrating a causal role for the insula in regulating compulsive drinking. Similarly, chemogenetic silencing of insula projections to the nucleus accumbens decreased alcohol self-administration and potentiated the subjective, or interoceptive, effects of alcohol in rats (*10-12*). Analogous behavioral studies in humans showed that heavy drinkers attempted to earn more alcohol drinks than light drinkers under threat of shock and exhibited greater connectivity between the insula and nucleus accumbens, a brain region involved in motivation and reward, when viewing shock-predictive alcohol cues (*3*). Higher connectivity between the insula and nucleus accumbens was also associated with compulsive alcohol use scores in these individuals. These studies demonstrate that the insula is involved in compulsive alcohol drinking and the interoceptive effects of alcohol, but specific molecular mechanisms within the insula that regulate these addiction-related behaviors have not been identified.

Cortical neurons that express the calcium-binding protein parvalbumin are fast-spiking, GABAergic interneurons that tightly regulate the firing of excitatory pyramidal projection neurons (*13, 14*). These neurons are critical for effective cognitive functioning and decision-making through their ability to maintain the proper balance between excitation and inhibition in the cortex (*15*). The majority of cortical parvalbumin-expressing interneurons are surrounded by specialized extracellular matrix structures known as perineuronal nets (PNNs), whose primary components are chondroitin sulfate proteoglycans and hyaluronan. PNNs play an important role in regulating the excitability of fast-spiking inhibitory interneurons by controlling their synaptic inputs (*16-18*). Synapses are located within the gaps of the lattice-like structure of PNNs, suggesting that PNNs can directly regulate synaptic architecture. Evidence that PNNs restrict synaptic plasticity come from studies demonstrating that PNNs form in an activity-dependent manner during the closure of critical periods of brain development in which plasticity is reduced (*19*). They are hypothesized to stabilize synapses during memory formation, leading to long-term memory storage, and prevent the formation of new memories (*20-22*). Recently, PNNs have gained prominence in the addiction research field through studies demonstrating that they regulate drug-associated memories that contribute to relapse (*23-27*).

Although evidence suggests that PNNs are likely involved in addiction to drugs such as cocaine and heroin, there are currently very few studies examining the role of PNNs in AUD. We previously found that repeated bouts of binge alcohol drinking in mice increased the intensity of PNNs in the insula (*28*), suggesting that alcohol induces adaptive changes in the extracellular matrix that could affect the functioning of parvalbumin-expressing neurons in this brain region. However, there are currently no studies demonstrating a functional role for PNNs in AUD. Because the insula regulates compulsive drinking, we hypothesized that manipulating insular PNNs in mice would alter alcohol drinking. We tested this hypothesis by measuring compulsive drinking in mice following digestion of PNNs in the insula. Our findings demonstrate that insular PNNs regulate aversion-resistant alcohol consumption and the activity of the insula during quinine-adulterated alcohol drinking. The implications are that alcohol-induced maladaptive increases in PNNs in the insula mediate the transition from more controlled to compulsive drinking and that interfering with PNNs in this brain region could be a means to restore control over compulsive drinking.

## Results

### Digestion of PNNs renders mice sensitive to quinine-adulterated ethanol

Disruption of PNN structure was achieved by using the enzyme chondroitinase ABC (ChABC), which digests the chondroitin sulfate chains on the proteoglycan constituents of PNNs (*29*). To disrupt PNNs in the insula and test the behavioral consequences, ChABC or phosphate-buffered saline (PBS, as a control) was microinjected directly into the anterior insula and mice were allowed to recover for 3 days prior to testing in a two-bottle choice test of ethanol consumption for 24 hours over the course of 4 days (**Fig. 1A**). The inclusion of bitter-tasting quinine in the ethanol solution effectively models aversion-resistant drinking in rodents (*9*), therefore mice were divided into two groups: one group received a choice between 15% ethanol *vs.* water to measure ethanol drinking and the other group received a choice between 15% ethanol containing 100 μm quinine (EQ) *vs.* water to measure aversion-resistant ethanol drinking. We found significant main effects of ChABC treatment and ethanol solution as well as a significant treatment by ethanol solution interaction in ethanol consumption (**Fig. 1B**, ChABC treatment: *F*_1, 124_ = 16.26, *p* < 0.0001; ethanol solution: *F*_1, 124_ = 36.34, *p* < 0.0001; interaction: F1, 124 = 5, *p* = 0.027). Mice treated with ChABC drank significantly less EQ solution compared with all of the other groups (*p* < 0.0001). Decreased EQ drinking in the ChABC group was evident on the first testing day and was maintained throughout the 4 days of drinking. Importantly, ChABC digestion of PNNs in the insula had no effect on ethanol drinking when quinine was not included in the ethanol solution, indicating a specific role for insular PNNs in aversion-resistant drinking as opposed to drinking under neutral circumstances. Mice treated with ChABC also had less preference for the EQ solution compared with all of the other groups, as measured by the ratio of the volume of EQ over total volume of fluid consumed (**Fig. 1C**, ChABC treatment, *F*_1, 124_ = 19.83, *p* < 0.0001; ethanol solution: *F*_1, 124_ = 19.13, *p* < 0.0001; interaction, *F*_1, 124_ = 7.86, *p* = 0.0059; post-hoc multiple comparisons test, *p* < 0.0001 for ChABC-treated mice drinking EQ compared with all groups). ChABC treatment did not alter total fluid consumption (**Fig. 1D**), demonstrating that digestion of PNNs did not adversely affect overall drinking behavior. Immediately after the drinking test, we confirmed PNN digestion in each mouse by staining PNNs in coronal brain sections throughout the insula using fluorescently-labeled *Wisteria floribunda* agglutinin (WFA), a plant protein that binds to chondroitin sulfate glycosaminoglycans in PNNs (*30*). PNNs were eliminated throughout the anterior portion of the insula on day 7 after behavioral testing (**Fig. 1E-F**). We also verified in a separate group of mice that PNNs were digested on day 4 when the drinking test began (**Fig. 1F**). These results indicate that PNNs in the insula are important for aversion-resistant drinking and that disrupting these structures can render mice more sensitive to quinine-adulterated ethanol.

**Fig. 1.**
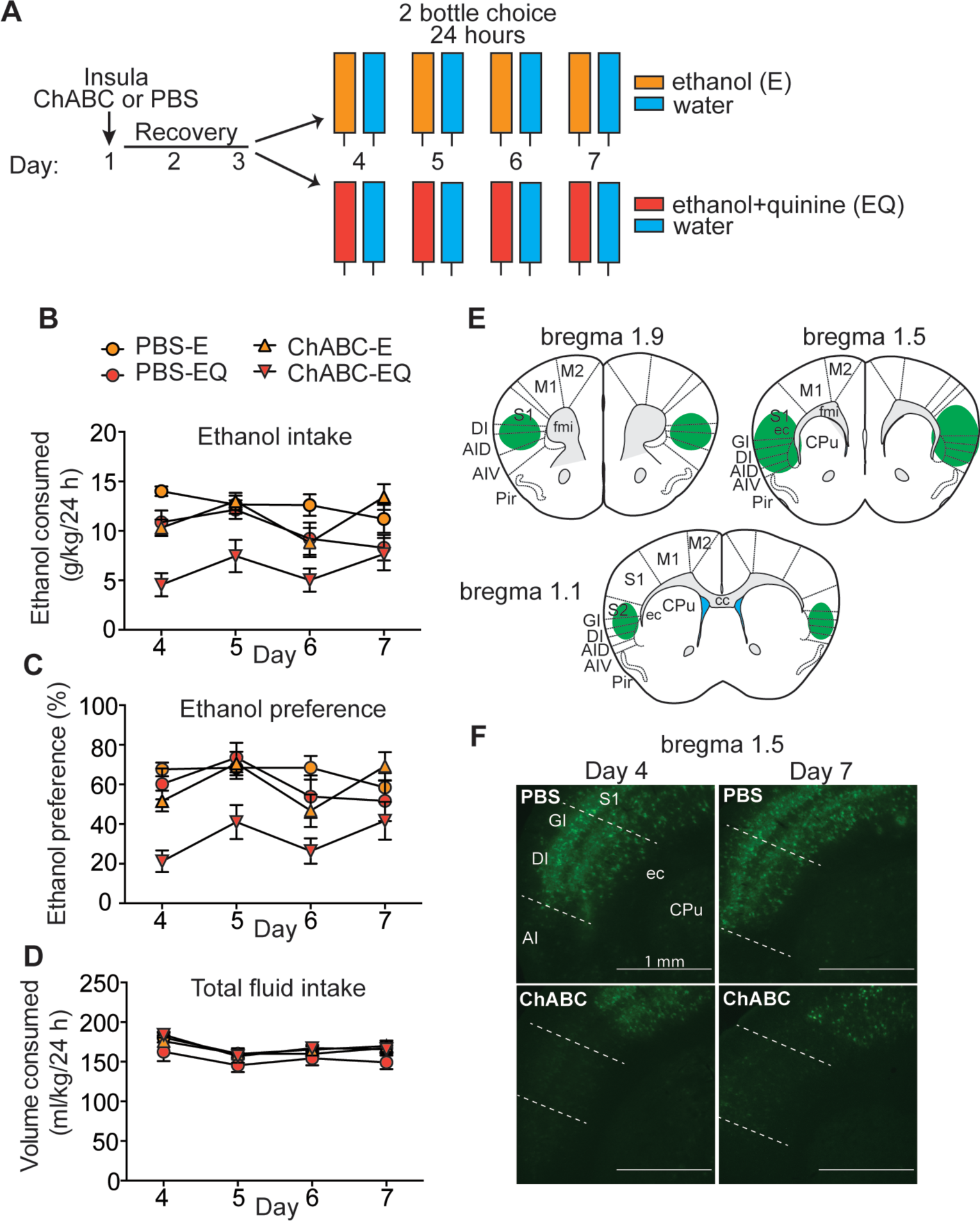
Digestion of perineuronal nets (PNNs) in the insula renders mice sensitive to quinine-adulterated ethanol. (**A**) Experimental design. Mice (n = 7-8 per group) were injected with either chondroitinase ABC (ChABC) or phosphate buffered saline (PBS) in the insula on day 1, recovered for 3 days, and then divided into 2 groups and given a two-bottle choice test for consumption of an ethanol solution (E) vs. water or ethanol plus quinine solution (EQ) vs. water for 24 hours over 4 days. (**B**) Daily ethanol consumption in g ethanol per kg body weight. (**C**) Daily ethanol preference (%), calculated as volume of ethanol solution consumed divided by total volume of fluid consumed. (**D**) Daily total fluid intake in ml per kg body weight. (**E**) Representation of injection site at 1.5 mm anterior to bregma and extent of PNN digestion in the insula (green areas) based on a mouse brain atlas. (**F**) Representative images of fluorescent *Wisteria floribunda* agglutinin binding to PNNs on coronal brain sections containing the insula on days 4 and 7 after injection of ChABC or PBS. Scalebar 1 mm; Abbreviations: CPu, caudate putamen; GI, granular insular cortex; DI, dysgranular insular cortex; AI, agranular insular cortex; S1, primary somatosensory cortex; ec, external capsule.

### Digestion of PNNs in the motor cortex does not affect ethanol drinking

The motor cortex is functionally distinct from the insula but contains a similar density and distribution of PNNs as the insula (*28*). To determine if the effect of PNN digestion on aversion-resistant drinking is specific to the insula, mice were injected with ChABC or PBS in the motor cortex and tested for ethanol drinking using the same protocol as above, except that drinking was tested for 1 day instead of 4 days, since the effect of insular PNN digestion on drinking was observed on the first day of testing (**Fig. 2A**). Mice with PNN digestion in the motor cortex did not significantly differ from controls in ethanol or EQ drinking or preference, nor did they differ in total fluid consumption (**Fig. 2B-D,** no main effects of ChABC treatment, ethanol solution, or interaction). PNN digestion was confirmed in the motor cortex (**Fig. 2E-F**), indicating that PNNs were completely digested by day 4 when the drinking test began. These results demonstrate that PNNs in the motor cortex do not regulate drinking of either ethanol or ethanol adulterated with quinine, and suggest that PNNs in the insula play a specific role in aversion-resistant drinking.

**Fig. 2.**
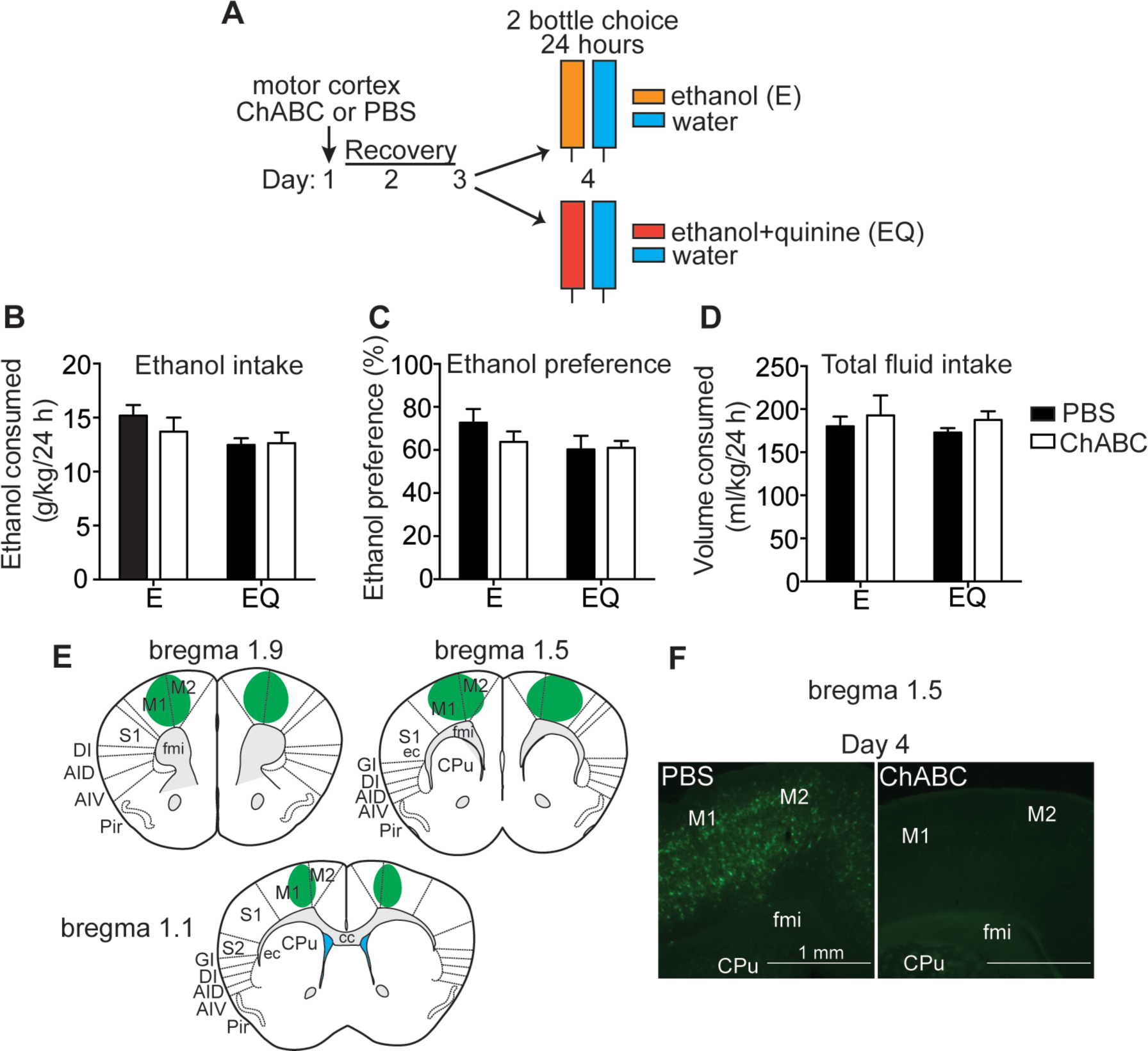
Digestion of perineuronal nets (PNNs) in the motor cortex does not affect ethanol drinking. (**A**) Experimental design. Mice (n= 6 per group) were injected with either chondroitinase ABC (ChABC) or phosphate buffered saline (PBS) in the motor cortex on day 1, recovered for 3 days, and then divided into 2 groups and given a 2-bottle choice test for consumption of an ethanol solution (E) vs. water or ethanol plus quinine solution (EQ) vs. water for 24 hours. (**B**) Daily ethanol consumption in g ethanol per kg body weight. (**C**) Daily ethanol preference (%), calculated as volume of ethanol solution consumed divided by total volume of fluid consumed. (**D**) Daily total fluid intake in ml per kg body weight. (**E**) Representation of injection site in the motor cortex based on a mouse brain atlas showing injected site at 1.5 mm anterior to bregma and spread of PNN digestion in green. (**F**) Representative images of fluorescent *Wisteria floribunda* agglutinin (WFA) binding to PNNs on coronal brain sections containing the motor cortex on day 4 after injection of ChABC or PBS. Scalebar, 1 mm. Abbreviations: CPu, caudate putamen; M1, primary motor cortex; M2, secondary motor cortex; fmi, forceps minor of the corpus callosum.

### Digestion of PNNs in the insula does not affect consumption of sucrose or quinine alone

The insula contains the gustatory cortex, a region that is critical for taste processing (*31*). To rule out the possibility that the effect of ChABC digestion on drinking EQ was simply due to increased sensitivity to bitter taste, mice were injected in the insula with ChABC or PBS and tested for two-bottle choice consumption of a 100 μm quinine solution *vs.* water for 24 hours (**Fig. 3A**). Digestion of PNNs in the insula had no effect on quinine drinking or preference relative to water (**Fig. 3B-C**), indicating that the effect of disrupting PNNs on aversion-resistant alcohol drinking is not simply due to increased sensitivity to the bitter taste of quinine. As an additional control, we also tested mice treated with ChABC for 2% sucrose drinking to determine if digesting PNNs in the insula altered sweet taste sensitivity. Sucrose drinking and preference were not altered by ChABC treatment (**Fig. 3D-E**), further demonstrating that digesting PNNs in the insula likely does not affect taste perception.

**Fig. 3.**
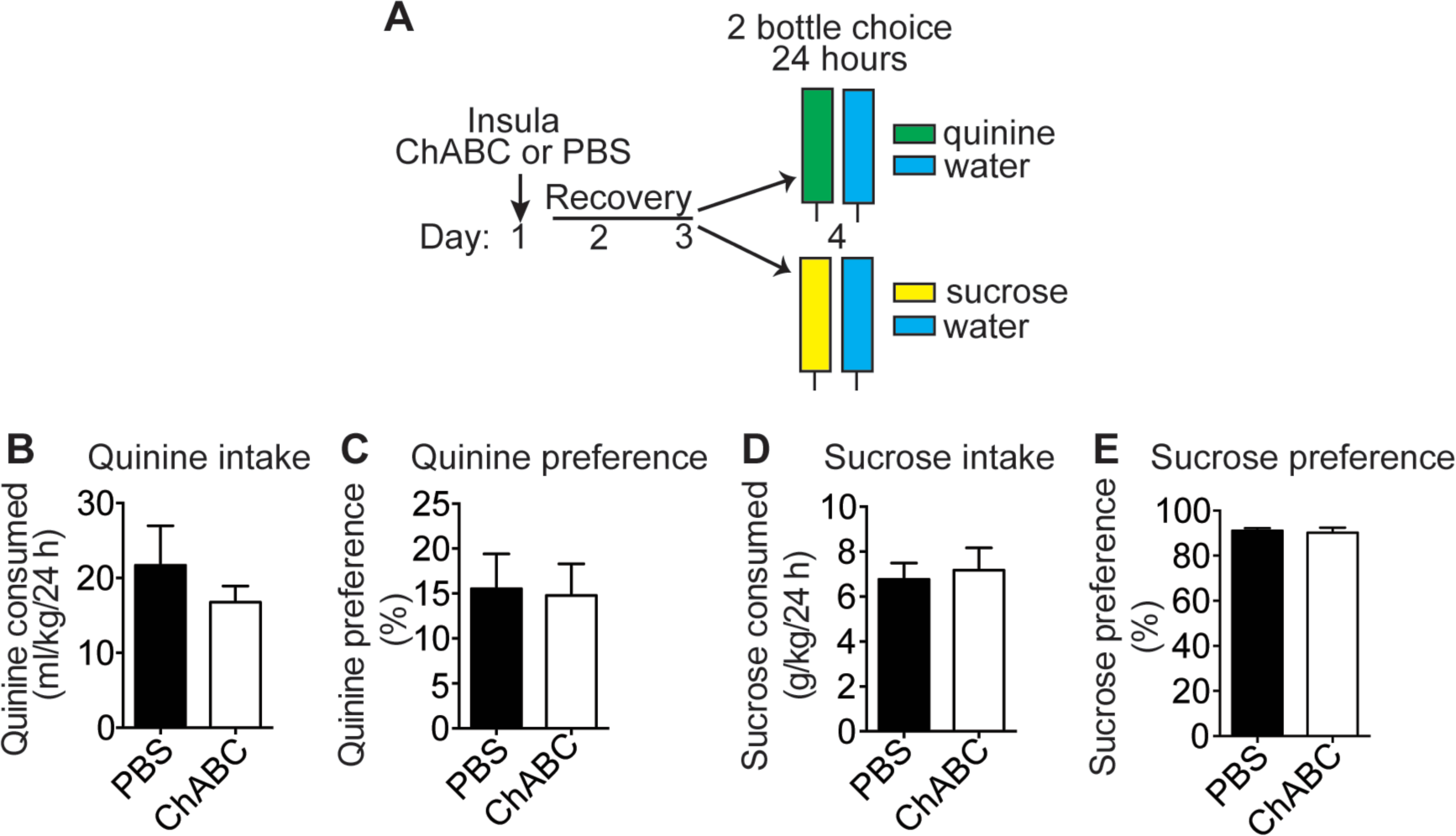
Digestion of perineuronal nets (PNNs) in the insula does not affect sucrose or quinine drinking. (**A**) Experimental design. Mice (n=6 per group) were injected with either chondroitinase ABC (ChABC) or phosphate buffered saline (PBS) in the insula on day 1, recovered for 2 days, and then were divided into 2 groups and given a two-bottle choice for 24 hours between a sucrose solution vs. water or a quinine solution vs. water. (**B**) Quinine solution consumed, in ml per kg body weight. (**C**) Percent quinine preference, calculated as volume of quinine solution consumed divided by volume of total fluid consumed. (**D**) Sucrose solution consumed, in g sucrose per kg body weight. (**E**) Percent sucrose preference, calculated as volume of sucrose solution consumed divided by volume of total fluid consumed.

### C-fos expression is induced in the insula of mice drinking quinine-adulterated ethanol

We hypothesized that during the initial stages of drinking a quinine-adulterated ethanol solution that the insula would become engaged because of its role in decision-making under conditions of conflict or risk (*5*). To test this, naïve mice were given a two-bottle choice test for 2 hours to measure expression of c-fos as a surrogate for neural activation. Drinking was conducted for 2 hours instead of 24 hours in order to capture peak c-fos expression during the drinking session. Mice were divided into four groups: control mice received water in both bottles, while the other three groups received either ethanol, quinine, or EQ in one of the bottles (**Fig. 4A**). During the 2-hour drinking session, mice that received ethanol or EQ drank comparable volumes of either ethanol or the EQ solution as the water drinking controls, whereas mice that received the quinine solution consumed significantly less of this solution compared with each of the other groups (**Fig. 4C**; significant effect of solution, *F*_3, 24_ = 5.1, *p* = 0.0070; post-hoc multiple comparisons test, *p* < 0.05 when comparing quinine to each of the other groups). Mice drank 2.8 g/kg and 2.6 g/kg ethanol from the ethanol or EQ bottle, respectively, during the 2-hour period. Mice also had similar preferences for water, ethanol, or EQ, but showed decreased preference for the quinine solution (**Fig. 4D**; significant effect of solution, *F*_3, 24_ = 8.92, *p* = 0.0004; post hoc multiple comparisons test, *p* < 0.01 when comparing quinine to each of the other groups), demonstrating that the quinine-only solution was clearly aversive. Total fluid consumption was equivalent in all four groups, however, indicating that the mice given quinine in one bottle were not merely consuming less overall fluid during the 2-hour session (**Fig. 4E**). Immediately after the 2-hour drinking session, brains were processed for immunohistochemistry using a c-fos antibody. Strikingly, the number of c-fos positive cells was significantly greater in mice that drank the EQ solution compared with all of the other groups (**Fig. 4F-G**; significant effect of solution, *F*_3, 103_ = 14.6, *p* < 0.0001; p < 0.001 by post-hoc multiple comparisons test when comparing EQ to each of the other groups). Expression of c-fos did not differ between mice drinking either ethanol- or quinine-only solutions compared with water (*p* = 0.074 for ethanol vs. water group and *p* = 0.34 for quinine vs. water group). These results suggest that the insula is not engaged during consumption of ethanol or bitter tastant alone, but instead is activated during drinking of quinine-adulterated ethanol, further supporting its role in the decision to continue drinking under conflict.

**Fig. 4.**
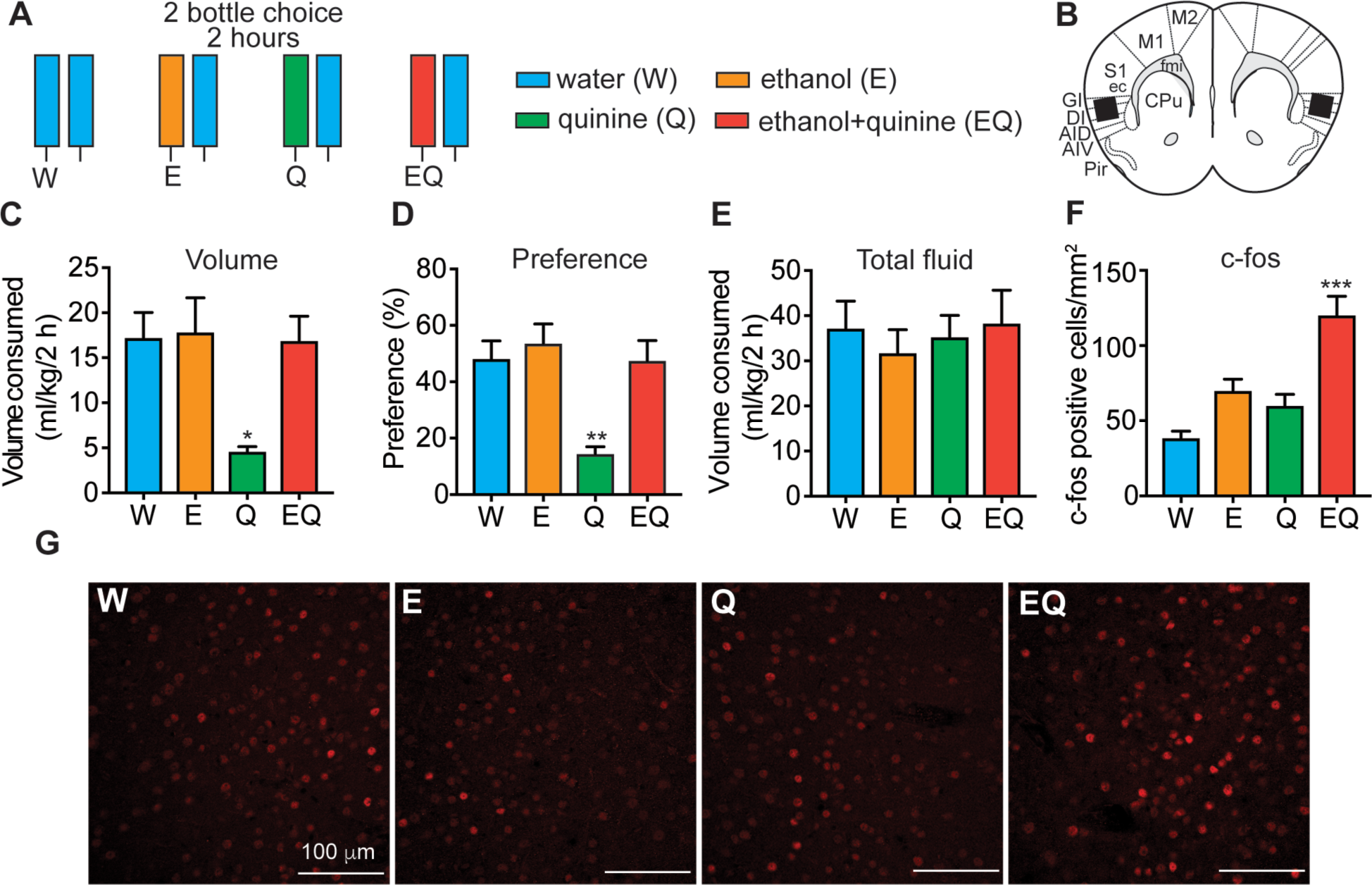
c-fos is induced in the insula of mice drinking quinine-adulterated ethanol. (**A**) Experimental design. Experimentally naïve mice (n=6) were given a 2-bottle choice test for 2 h followed by perfusion and collection of brain tissue for immunohistochemistry with a c-fos antibody. Mice were divided into 4 groups and given different 2-bottle choice tests: water (W) vs. W, ethanol (E) vs. W, quinine (Q) vs. W, and ethanol plus quinine (EQ) vs. W. (**B**) Diagram of area in which c-fos counts were obtained based on the mouse brain atlas. Black box shows area where counts were taken. (**C**) Volume of one bottle consumption of W, E, Q, and EQ in ml per kg body weight. \**p* < 0.05 by Tukey’s post-hoc multiple comparisons test when comparing Q to all other groups. (**D**) Percent preference of each solution (W, E, Q, or EQ), calculated as volume of solution consumed divided by total volume consumed. \*\**p* < 0.01 by Tukey’s post-hoc multiple comparisons test when comparing Q to all other groups. (**E**) Total fluid consumed in ml per kg body weight. (**F**) Number of c-fos positive cells in the insula of mice drinking W, E, Q, and EQ. \*\*\**p* < 0.001 when by Tukey’s post-hoc multiple comparisons test when comparing EQ to all other groups. Data are presented as average c-fos counts from 6 mice per group and 4-5 sections per mouse. (**G**) Representative images showing c-fos immunoreactivity in the insula 2 h after consuming W, E, Q, and EQ. Scalebars, 100 μm.

### c-fos is induced in glutamatergic neurons in the insula of mice drinking quinine-adulterated ethanol

In the insula, PNNs primarily surround parvalbumin-expressing GABAergic interneurons and regulate their excitability (*17, 18*). We speculated that c-fos might be induced in these neurons during EQ drinking because digestion of PNNs in the insula alters consumption of EQ. We therefore examined which type(s) of neurons in the insula express c-fos during EQ drinking by performing dual fluorescent immunohistochemistry on brain sections containing the insula from mice that drank EQ for 2 hours. Brain sections were fluorescently labeled with antibodies to c-fos and WFA (for PNN-enclosed cells), parvalbumin, or CaMKII-a (for glutamatergic neurons). Unexpectedly, c-fos expression was almost entirely localized to CaMKII-a-expressing neurons (**Fig. 5**), not parvalbumin-expressing neurons or neurons with PNNs. These data indicate that the insular neurons activated during EQ drinking are primarily glutamateric.

**Fig. 5.**
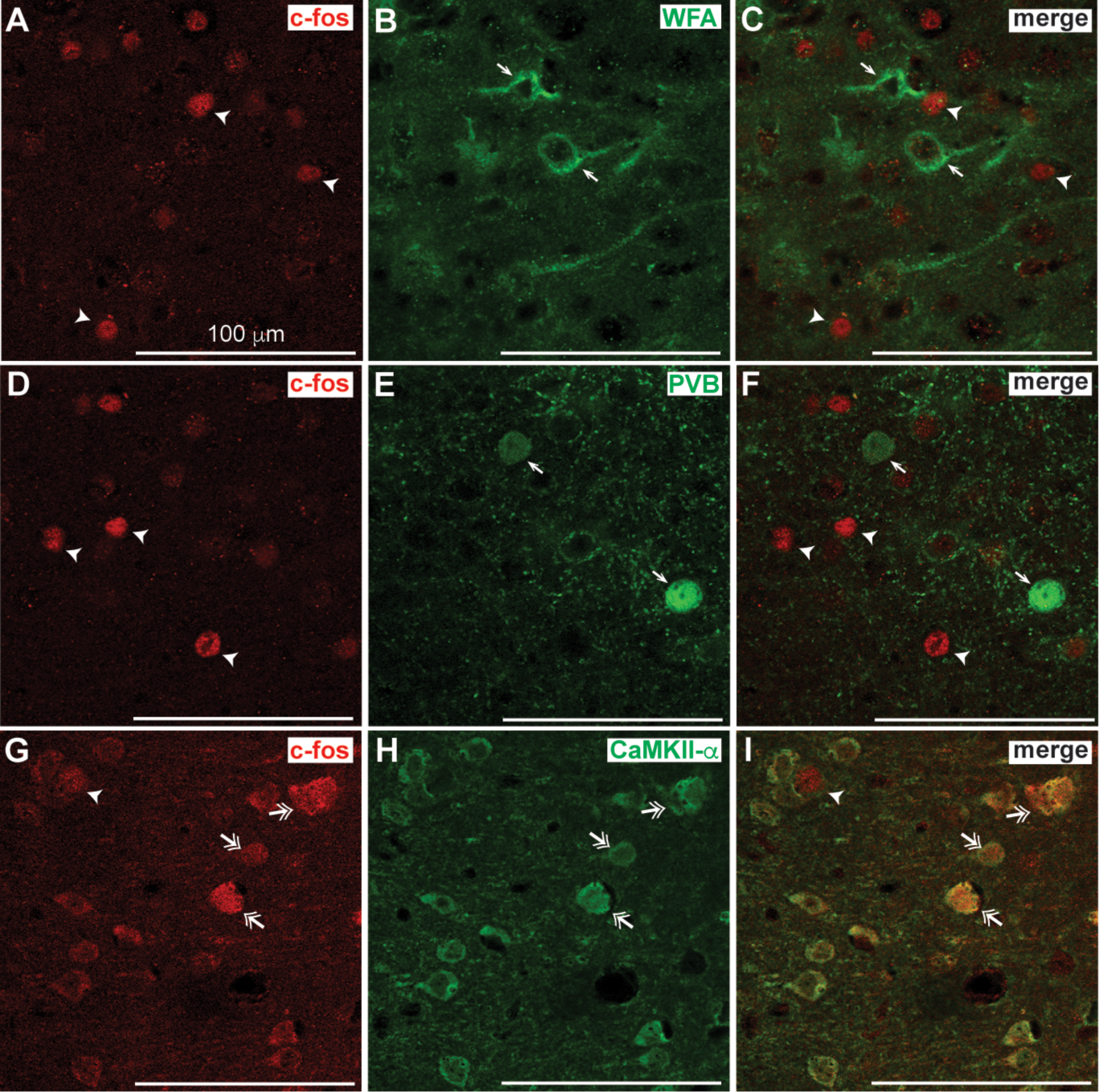
C-fos expression is induced in glutamatergic neurons in the insula of mice drinking quinine-adulterated ethanol. Experimentally naïve mice (n=6) were given a 2-bottle choice test for ethanol plus quinine vs. water for 2 h followed by perfusion and collection of brain tissue for dual fluorescent immunohistochemistry with antibodies to c-fos and parvalbumin (PVB), calcium/calmodulin dependent protein kinase II alpha (CaMKII-a), or fluorescently-labeled *Wisteria floribunda* agglutinin (WFA). Shown are representative single-channel fluorescent images of c-fos (**A, D,** and **G**, in red) and (**B**) WFA, (**E**) PVB and (**H**) CaMKII-a (in green) followed by the merged images in (**C, F,** and **I**). Arrowheads indicate cells that are positive for cfos only, while single arrows indicate cells that are positive for WFA (**B**) or PVB (**E**) only. Double arrowheads indicate cells that are positive for both c-fos and CaMKII-a. Note the co-localization of c-fos with CaMKII-a but not PVB- or WFA-positive cells. Scalebar, 100 μm.

### c-fos induction is elevated during drinking of quinine-adulterated ethanol in mice lacking PNNs in the insula

We next reasoned that, although c-fos was induced in glutamatergic neurons in the insula during EQ drinking, digestion of PNNs might alter activation of these neurons during drinking of quinine-adulterated ethanol because of the ability of PNNs to regulate the excitability of fast-spiking interneurons which tightly control the firing of cortical pyramidal neurons (*14*). To test this, mice were injected with ChABC or PBS in the insula, recovered for 3 days, and then underwent a 2-hour drinking session (again, to capture peak c-fos expression) for either ethanol or EQ before measuring c-fos immunoreactivity in the insula (**Fig. 6A**). Ethanol intake (g/kg), preference, and total fluid consumed were not significantly different between groups after 2 hours of drinking, although there was a slight decrease in EQ drinking and preference in the ChABC-treated group (**Fig. 6B-C**), with a trend towards a ChABC treatment by ethanol solution interaction in g/kg intake (*F*_1, 20_ = 3.5, *p* = 0.076) and a significant interaction in preference (*F*_1, 20_ = 4.91, *p* = 0.039), but none of the comparisons was significant after multiple comparisons testing. This was not entirely unexpected even though ChABC-treatment resulted in decreased EQ drinking during a 24-hour session because of the fairly short length of the drinking session. However, the number of c-fos positive cells in the insula was significantly higher in mice drinking EQ (**Fig. 6E-F**; main effect of ethanol solution, *F* _1, 104_ = 25.07, *p* < 0.0001) and trended toward significance in mice treated with ChABC (main effect of treatment, *F* _1, 104_ = 3.46, *p* = 0.066). These results demonstrate that manipulating PNNs in the insula leads to greater activation of the insula during the initial drinking of quinine-adulterated ethanol.

**Fig. 6.**
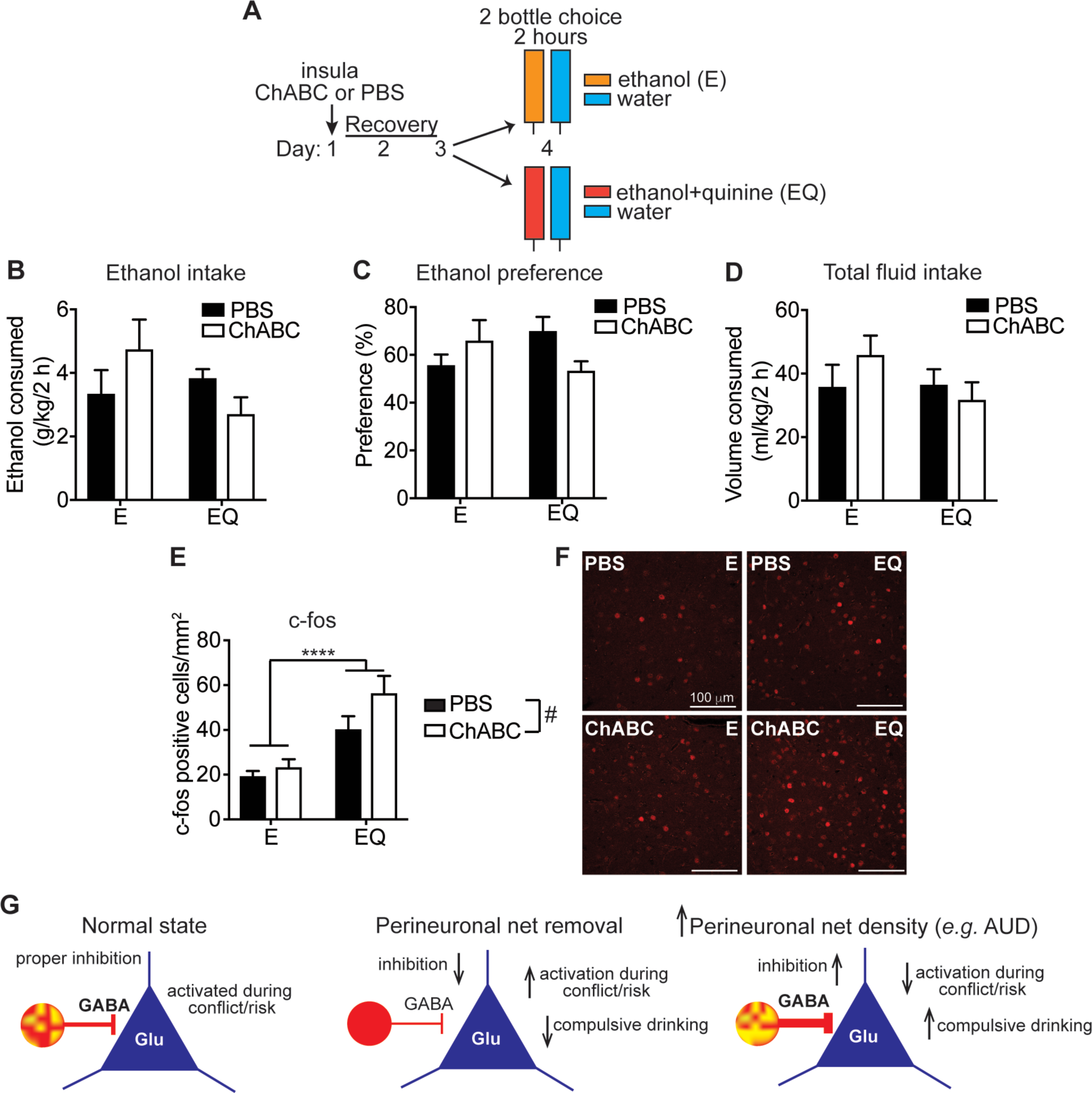
c-fos induction is elevated during drinking quinine adulterated ethanol in mice lacking perineuronal nets (PNNs) in the insula. (**A**) Experimental Design. Mice (n=6) were injected with either chondroitinase ABC (ChABC) or phosphate buffered saline (PBS) in the insula on day 1, recovered for 2 days, and then were divided into 2 groups and given a 2-bottle choice for 2 h between ethanol (E) vs. water or ethanol plus quinine (EQ) vs. water followed by perfusion and collection of brain tissue for fluorescent immunohistochemistry with an antibody to c-fos. (**B**) Amount of E or EQ consumed in a 2 h period, expressed as g per kg body weight. (**C**) Percent preference for E or EQ, calculated as volume of E or EQ consumed divided by total fluid consumption. (**D**) Total fluid consumed in 2 h, expressed as ml per kg body weight. (**E**) Number of c-fos positive cells in the insula. \*\*\*\**p* < 0.0001, main effect of ethanol solution by two-way ANOVA; ^#^*p* = 0.066, main effect of ChABC by two-way ANOVA. Data are presented as average c-fos counts from 6 mice per group and 4-5 sections per mouse. (**F**) Representative images of c-fos immunoreactivity in the insula. Scalebars, 100 μm. (**G**) Model of inhibitory control of pyramidal neurons (blue triangles) by parvalbumin-expressing GABA interneurons (red circles) under normal conditions, after PNN removal by ChABC treatment, and after chronic alcohol drinking or alcohol use disorder (AUD) and hypothesized relationship to compulsive drinking. Density of PNNs is indicated as intensity of yellow staining on red parvalbumin neurons.

## Discussion

The insula is an integrator of sensory, emotional, motivational, and cognitive functions and has emerged as a key brain region involved in addiction to several drugs of abuse. Here, we provide evidence that degradation of PNNs in the insula decreases quinine-adulterated ethanol drinking, suggesting these extracellular matrix structures are important in promoting compulsive alcohol consumption. Our results also indicate that PNNs control the activity of the insula early during drinking of quinine-adulterated alcohol when animals must decide whether to continue drinking alcohol despite the addition of a bitter tastant. The insula is activated during drinking of quinine-adulterated ethanol and this activation is augmented when PNNs are disrupted. Subsequently, mice drink less quinine-adulterated ethanol. Strikingly, we did not observe significant induction of c-fos when mice were drinking solutions containing only ethanol or quinine, nor did PNN digestion affect drinking of the ethanol- or quinine-only solutions. The combination of ethanol and quinine likely contains both rewarding and aversive elements, thus deciding whether to drink EQ likely presents a conflict to the animal. The implication of our results is that when there is no conflict, the insula is not activated and therefore does not necessarily play a significant role in alcohol drinking under risk-free circumstances.

One interpretation of our data is that greater insula activity facilitates better decision-making under conditions of conflict or risk. A study in human subjects with AUD found that greater anterior insula activation is correlated with decreased risk-taking (*8*). Brain imaging studies have also shown that the size of the anterior insula is decreased in alcohol-dependent individuals compared with healthy controls (*7, 32*) and that a smaller anterior insula predicts increased compulsivity (*7*). The deficits in insula volume could be caused by chronic alcohol exposure, or could be a predisposing factor for AUD. In rats, chemogenetic silencing of the anterior insula increases alcohol self-administration (*11*). Together, these studies indicate that individuals with AUD are unable to appropriately engage the insula during risky decision-making because of structural deficits in the insula.

Notably, intoxicating doses of alcohol also reduce the activity of the insula, increase risk-taking behavior (*33, 34*), and dampen the response of the anterior insula to notification of a negative outcome after risk-taking (*33*). Decreased insula activity in response to alcohol intoxication is blocked by a GABA_A_ receptor antagonist, demonstrating that inhibitory neurotransmission is responsible for dampening insula activity during intoxication (*34*). Silencing the insula also potentiates the interoceptive effects of alcohol (*10*). We propose that greater activation of the anterior insula during risky alcohol drinking, in which there is high probability of a negative outcome if drinking continues, results in a decision to cease drinking. Further, this suggests that individuals with AUD drink compulsively because they are unable to appropriately engage the insula while consuming alcohol and thus continue to drink despite risk. This hypothesis is consistent with decreased anterior insula volume observed in heavy drinkers, which would prohibit optimal activation of the insula while drinking when negative consequences are likely. Compulsive drinkers might also be more sensitive to alcohol-induced inactivation of the insula.

However, these results are in apparent contradiction with other studies demonstrating that heavy drinkers show greater connectivity between the anterior insula and nucleus accumbens than light drinkers when viewing alcohol cues that predict electric shock and attempt to earn more alcohol during the high-threat conditions (*3*). Similarly, Seif *et al* showed in rats that inhibition of insular inputs to the nucleus accumbens decreases aversion-resistant drinking, using an ethanol and quinine drinking protocol similar to the one used in this study, and Pushparaj and Foll found that pharmacological silencing of the posterior granular insula in rats decreases alcohol self-administration (*4, 35*). These discrepancies with our findings might be reconciled by differences in the activation of specific insular circuits. The insula sends projections to brain regions involved in both reward and aversion (*36*), including the amygdala, and activation of different circuits during decision-making likely differentially guide behavior. Indeed, chemogenetic silencing of insula to nucleus accumbens afferents decreases ethanol self-administration, yet silencing of insula itself increases ethanol self-administration (*11*), implicating different circuits in mediating these opposing behavioral responses. One limitation of our study is that we did not measure the contribution of individual circuits when examining c-fos expression. We measured c-fos in cortical layers II/III of the rostral granular and dysgranular insular cortex, where PNNs are abundant. Tracing studies in rats indicate that these regions of the insula send efferents to limbic regions such as the amygdala and lateral hypothalamus (*36*) and to the agranular insular cortex (*37*), whereas the anterior agranular insular cortex projects to the nucleus accumbens (*38*). More studies are needed to dissect the role of specific insular subregions and circuits in guiding aversion-resistant versus aversion-sensitive alcohol drinking.

Cortical PNNs primarily surround fast-spiking parvalbumin-positive interneurons that are critical for the maintenance of a proper excitatory/inhibitory balance. The increased activation of glutamatergic neurons after PNN removal in the insula during quinine-adulterated ethanol drinking suggests that PNN digestion disinhibits pyramidal projection neurons, resulting in increased cortical excitability (**Fig. 6G**). In support of this hypothesis, several studies have found that digestion of PNNs with ChABC decreases firing of inhibitory interneurons and their input onto pyramidal cells (*17, 18, 25, 39*). *In vivo* recordings of cortical neurons after ChABC digestion also show decreased inhibitory activity and a disruption in the excitatory/inhibitory balance (*40*). Our results suggest that strong inhibitory control over insula principal neurons is required for compulsive-like ethanol drinking.

Our previous work showed that the density of PNNs around neurons in the insula increased after repeated bouts of binge drinking in mice (*28*). Increased thickness of PNNs is predicted to facilitate rapid firing of parvalbumin neurons, thus increasing inhibitory control over excitatory projection neurons. As a result, it may be more difficult to activate the insula after chronic alcohol exposure when PNNs are more pronounced. Together with the results presented here, these data suggest that increased PNNs and the resulting GABAergic inhibition in the insula after repeated bouts of binge drinking plays a role in the transition from social, non-compulsive drinking to compulsive alcohol abuse.

Our results suggest that disrupting PNNs in the insula and restoring normal excitatory/inhibitory balance in the insula may be a novel therapeutic approach to treating compulsive drinking. Injection of an extracellular matrix-degrading enzyme such as ChABC into the insula is not a viable therapeutic option, given the invasive and temporary nature of this manipulation, but a pharmacotherapeutic aimed at reducing PNNs might be possible. The extracellular matrix has gained prominence in recent years as yet another mechanism involved in the regulation of synaptic plasticity and neuronal activity. Our results provide further evidence that PNNs are important for addiction but play a specific role in the insula in controlling compulsive drinking. Degrading PNNs, decreasing interneuron excitability, or increasing excitatory neurotransmission in the insular cortex may restore healthy insula function in individuals with AUD, leading to better decision-making under conditions of risk or conflict.

## Materials and Methods

### Experimental design

Experimental groups, timing of experimental treatments, and behavioral testing in mice are shown in panel A of Figs. 1-4 and 6.

### Animals

Male C57BL/6J mice were purchased from The Jackson Laboratory at the age of 8 weeks (Bar Harbor, ME, USA). Mice were individually housed in a temperature- and humidity-controlled room with a 12-hour reversed light/dark cycle (lights off at 10 am) for 2 weeks prior to beginning experiments. Food and water were available *ad libitum*. All procedures with mice were conducted according to the National Institutes of Health *Guide for the Care and Use of Laboratory Animals* and approved by the UIC Animal Care and Use Committee.

### Chemicals and antibodies

Biotinylated WFA and Dylight 488-conjugated streptavidin were purchased from Vector Laboratories (WFAB-1355 and SA-5488, Burlingame, CA, USA). Mouse anti-calcium/calmodulin-dependent protein kinase II alpha (CaMKII-α) was purchased from Millipore (05-532, Billerica, MA, USA). Mouse anti-parvalbumin (PV) was purchased from synaptic systems (195011, Goettingen, Germany). Rabbit anti-c-Fos was purchased from Santa Cruz Biotechnology (SC-52, Dallas, TX, USA). Corresponding Alexa Flour 488- and Alexa Fluor 594-conjugated secondary antibodies were from Jackson ImmunoResearch (715-545-151 and 711-585-152, West Grove, PA, USA). ChABC (C-3667) and quinine were purchased from Sigma-Aldrich (St. Louis, MO, USA).

### ChABC treatment

After acclimating to a reversed light/dark cycle for 2 weeks, mice were anesthetized with ketamine (100 mg/kg) and xylazine (8 mg/kg), placed on a stereotaxic apparatus, and the skull prepared for intracranial injections of ChABC or PBS using 33 gauge stainless steel hypodermic tubing connected to an infusion pump. Mice were injected bilaterally with 0.5 μl ChABC (50 units/ml) or PBS into the insula or motor cortex. Coordinates for bilateral infusion were: insula, anterior/posterior, +1.5 mm from bregma; medial/lateral, +/− 2.9 mm from midline; and dorsal/ventral −2.6 mm from the top of the skull; motor cortex, anterior/posterior, +1.5 mm from bregma; medial/lateral, +/− 1.5 mm from midline; and dorsal/ventral −1.5 mm from the top of the skull. Mice recovered for 3 days prior to testing drinking.

### Two-bottle choice drinking tests

Water and 15% ethanol in water (E), or water and 15% ethanol with 100 μM quinine in water (EQ) were provided to mice in 10 ml clear polystyrene serological pipets (Fisher Scientific) truncated at the end to accommodate connection to a 2.5-inch stainless steel ball-bearing sipper tube (Ancare Corp, Bellmore, NY, USA). For ethanol drinking measured in **Fig. 1**, mice were given access to the 2 tubes in the home cage for 24 hours per day for 4 days, with consumption measured every day. Tube placements were alternated daily to avoid the confound of preference for a particular side. For ethanol drinking measured in **Fig. 2**, mice were given access to 2 tubes in the home cage as indicated above, but only for 1 day instead of 4 days. For the taste preference test in **Fig. 3**, mice were given access to either water and 2% sucrose in water (w/v), or water and 100 μM quinine in water. For the measurement of c-fos in **Figs. 4-6**, mice were divided into four groups and given one of the following 2-bottle choice tests: water/water, E/water, quinine (Q)/water, or EQ/water for 2 hours during the dark cycle, 3 hours after lights off.

### Immunohistochemistry

Immediately after the drinking tests, mice were euthanized using a lethal dose of a commercial euthanasia solution containing 390 mg pentobarbital/ml (Somnasol) and transcardially perfused with ice-cold phosphate buffered saline (PBS), followed by 4% paraformaldehyde (PFA). Brains were dissected and post-fixed in PFA overnight and cryoprotected in 30% sucrose for 48 hours. Every 4^th^ section of 50 μm serial coronal sections were collected from the area of the brain containing the insula, spanning +2.0 to +1.0 mm anterior to bregma. Sections were blocked in 5% normal donkey serum (017-000121, Jackson Immunoresearch) for 1 hour before incubation with c-Fos (1:500) for 48 hours at 4°C, followed by incubation with Alexa Fluor 594-conjugated donkey anti-rabbit (1:500) for 1 hour. After rinsing in PBS and re-blocking, sections were treated with WFA (1:2000), CaMKII-α (1:1000) or PV (1:1000) overnight at 4°C, then incubated with 488-conjugated secondary antibodies for 1 hour. Sections stained with WFA were blocked with carbo-free blocking solution instead of donkey serum (SP-5040-125, Vector Laboratories). For CaMKII-α staining, sections were heated in 10 mM citrate buffer (pH 8.5) at 80°C for 30 min before performing the immunostaining protocol above. Images were captured using a Zeiss LSM710 confocal microscope (Carl Zeiss, Thornwood, NY, USA) for cell analysis. An Evos FL microscope (Thermo Fisher Scientific, Waltham, MA USA) was used for visualization of ChABC digestion. C-fos counts were obtained from 6 mice per group and 4-5 sections per mouse, with numbers of c-fos positive cells averaged across all mice and sections. Cells were analyzed in cortical layers II/III of the granular and dysgranular insula, where PNNs predominate. Cell counting was performed manually using ImageJ software (National Institutes of Health, Bethesda, MD USA) and a threshold was applied during image analysis to normalize background staining.

### Statistical analysis

Data are presented as the mean ± SEM. Statistical testing was performed using Prism software (version 7, GraphPad, La Jolla, CA). For the ethanol drinking tests in **Fig. 1**, a three-way ANOVA was first performed for variables of day, treatment, and ethanol solution. There was a significant interaction between treatment and ethanol solution, but no significant three-way interaction. Therefore, data was collapsed across days and a two-way ANOVA was performed for effects of treatment and ethanol solution, followed by post-hoc Tukey’s multiple comparisons tests. For the drinking tests in **Figs. 2** and **6**, a two-way ANOVA was performed for effects of treatment and ethanol solution. A student’s t-test was performed for the data in **Fig. 3**, and a oneway ANOVA was performed for the data in **Fig. 4** followed by Tukey’s post hoc multiple comparisons tests. A *p* value of less than 0.05 was accepted as statistically significant.

## Acknowledgements

We would like to thank Dr. Elizabeth Glover and Dr. Amynah Pradhan for helpful comments on this manuscript. This work was supported by the National Institute on Alcohol Abuse and Alcoholism of the National Institutes of Health (INIA consortium grant U01 AA020912 and the Center for Alcohol Research in Epigenetics grant P50 AA022538). The content is solely the responsibility of the authors and does not necessarily represent the official views of the National Institute of Health.

## References

1. B. F. Grant et al., Epidemiology of DSM-5 Alcohol Use Disorder: Results From the National Epidemiologic Survey on Alcohol and Related Conditions III. JAMA Psychiatry 72, 757-766 (2015).

2. H. R. Kranzler, M. Soyka, Diagnosis and Pharmacotherapy of Alcohol Use Disorder: A Review. JAMA 320, 815-824 (2018).

3. E. N. Grodin et al., Neural Correlates of Compulsive Alcohol Seeking in Heavy Drinkers. Biol Psychiatry Cogn Neurosci Neuroimaging, (2018).

4. T. Seif et al., Cortical activation of accumbens hyperpolarization-active NMDARs mediates aversion-resistant alcohol intake. Nature neuroscience 16, 1094-1100 (2013).

5. N. Gogolla, The insular cortex. Curr Biol 27, R580-R586 (2017).

6. N. H. Naqvi, D. Rudrauf, H. Damasio, A. Bechara, Damage to the insula disrupts addiction to cigarette smoking. Science 315, 531-534 (2007).

7. E. N. Grodin, C. R. Cortes, P. A. Spagnolo, R. Momenan, Structural deficits in salience network regions are associated with increased impulsivity and compulsivity in alcohol dependence. Drug Alcohol Depend 179, 100-108 (2017).

8. E. D. Claus, K. E. Hutchison, Neural mechanisms of risk taking and relationships with hazardous drinking. Alcohol Clin Exp Res 36, 932-940 (2012).

9. F. W. Hopf, H. M. Lesscher, Rodent models for compulsive alcohol intake. Alcohol 48, 253-264 (2014).

10. A. A. Jaramillo et al., Functional role for suppression of the insular-striatal circuit in modulating interoceptive effects of alcohol. Addict Biol, (2017).

11. A. A. Jaramillo et al., Functional role for cortical-striatal circuitry in modulating alcohol self-administration. Neuropharmacology 130, 42-53 (2018).

12. A. A. Jaramillo, K. Van Voorhies, P. A. Randall, J. Besheer, Silencing the insular-striatal circuit decreases alcohol self-administration and increases sensitivity to alcohol. Behav Brain Res 348, 74-81 (2018).

13. D. Kvitsiani et al., Distinct behavioural and network correlates of two interneuron types in prefrontal cortex. Nature 498, 363-366 (2013).

14. N. Dehorter, N. Marichal, O. Marin, B. Berninger, Tuning neural circuits by turning the interneuron knob. Curr Opin Neurobiol 42, 144-151 (2017).

15. B. R. Ferguson, W. J. Gao, PV Interneurons: Critical Regulators of E/I Balance for Prefrontal Cortex-Dependent Behavior and Psychiatric Disorders. Front Neural Circuits 12, 37 (2018).

16. W. Hartig, K. Brauer, V. Bigl, G. Bruckner, Chondroitin sulfate proteoglycan-immunoreactivity of lectin-labeled perineuronal nets around parvalbumin-containing neurons. Brain research 635, 307-311 (1994).

17. T. S. Balmer, Perineuronal Nets Enhance the Excitability of Fast-Spiking Neurons. eNeuro 3, (2016).

18. P. Chu et al., The Impact of Perineuronal Net Digestion Using Chondroitinase ABC on the Intrinsic Physiology of Cortical Neurons. Neuroscience 388, 23-35 (2018).

19. T. Pizzorusso et al., Reactivation of ocular dominance plasticity in the adult visual cortex. Science 298, 1248-1251 (2002).

20. N. Gogolla, P. Caroni, A. Luthi, C. Herry, Perineuronal nets protect fear memories from erasure. Science 325, 1258-1261 (2009).

21. R. Y. Tsien, Very long-term memories may be stored in the pattern of holes in the perineuronal net. Proc Natl Acad Sci U S A 110, 12456-12461 (2013).

22. E. H. Thompson et al., Removal of perineuronal nets disrupts recall of a remote fear memory. Proc Natl Acad Sci U S A 115, 607-612 (2018).

23. J. M. Blacktop, R. P. Todd, B. A. Sorg, Role of perineuronal nets in the anterior dorsal lateral hypothalamic area in the acquisition of cocaine-induced conditioned place preference and self-administration. Neuropharmacology 118, 124-136 (2017).

24. B. R. Lubbers et al., The Extracellular Matrix Protein Brevican Limits Time-Dependent Enhancement of Cocaine Conditioned Place Preference. Neuropsychopharmacology 41, 1907-1916 (2016).

25. M. Slaker et al., Removal of perineuronal nets in the medial prefrontal cortex impairs the acquisition and reconsolidation of a cocaine-induced conditioned place preference memory. J Neurosci 35, 4190-4202 (2015).

26. M. C. Van den Oever et al., Extracellular matrix plasticity and GABAergic inhibition of prefrontal cortex pyramidal cells facilitates relapse to heroin seeking. Neuropsychopharmacology 35, 2120-2133 (2010).

27. Y. X. Xue et al., Depletion of perineuronal nets in the amygdala to enhance the erasure of drug memories. J Neurosci 34, 6647-6658 (2014).

28. H. Chen, D. He, A. W. Lasek, Repeated Binge Drinking Increases Perineuronal Nets in the Insular Cortex. Alcohol Clin Exp Res 39, 1930-1938 (2015).

29. L. D. Moon, R. A. Asher, K. E. Rhodes, J. W. Fawcett, Regeneration of CNS axons back to their target following treatment of adult rat brain with chondroitinase ABC. Nature neuroscience 4, 465-466 (2001).

30. W. Hartig, K. Brauer, G. Bruckner, Wisteria floribunda agglutinin-labelled nets surround parvalbumin-containing neurons. Neuroreport 3, 869-872 (1992).

31. A. Maffei, M. Haley, A. Fontanini, Neural processing of gustatory information in insular circuits. Curr Opin Neurobiol 22, 709-716 (2012).

32. V. V. Senatorov et al., Reduced anterior insula, enlarged amygdala in alcoholism and associated depleted von Economo neurons. Brain 138, 69-79 (2015).

33. J. M. Gilman, A. R. Smith, V. A. Ramchandani, R. Momenan, D. W. Hommer, The effect of intravenous alcohol on the neural correlates of risky decision making in healthy social drinkers. Addict Biol 17, 465-478 (2012).

34. T. Tsurugizawa, A. Uematsu, H. Uneyama, K. Torii, The role of the GABAergic and dopaminergic systems in the brain response to an intragastric load of alcohol in conscious rats. Neuroscience 171, 451-460 (2010).

35. A. Pushparaj, B. Le Foll, Involvement of the caudal granular insular cortex in alcohol self-administration in rats. Behav Brain Res 293, 203-207 (2015).

36. G. V. Allen, C. B. Saper, K. M. Hurley, D. F. Cechetto, Organization of visceral and limbic connections in the insular cortex of the rat. J Comp Neurol 311, 1-16 (1991).

37. C. J. Shi, M. D. Cassell, Cortical, thalamic, and amygdaloid connections of the anterior and posterior insular cortices. J Comp Neurol 399, 440-468 (1998).

38. S. M. Reynolds, D. S. Zahm, Specificity in the projections of prefrontal and insular cortex to ventral striatopallidum and the extended amygdala. J Neurosci 25, 11757-11767 (2005).

39. B. P. Tewari et al., Perineuronal nets decrease membrane capacitance of peritumoral fast spiking interneurons in a model of epilepsy. Nat Commun 9, 4724 (2018).

40. K. K. Lensjo, M. E. Lepperod, G. Dick, T. Hafting, M. Fyhn, Removal of Perineuronal Nets Unlocks Juvenile Plasticity Through Network Mechanisms of Decreased Inhibition and Increased Gamma Activity. J Neurosci 37, 1269-1283 (2017).

